# Distinct resting-state connectomes for face and scene perception predict individual task performance

**DOI:** 10.1101/2025.07.09.663812

**Authors:** Orhan Soyuhos, Aurelia Scarpa, Daniel Baldauf

## Abstract

Face and scene perception rely on distinct neural networks centered on the Fusiform Face Area (FFA) and Parahippocampal Place Area (PPA). However, how these regions interact with broader brain networks remains unclear. Using resting-state fMRI and MEG data, we mapped the spatial and frequency-specific functional connectivity of the FFA and PPA. We found that the FFA showed predominant fMRI connectivity with lateral occipitotemporal, inferior temporal, and temporoparietal regions, while the PPA connected more strongly with ventral medial visual, posterior cingulate, and entorhinal-perirhinal areas. MEG analyses further revealed this network segregation was reflected in beta and gamma bands. Importantly, connectome-based predictive modeling showed that the strength of these intrinsic fMRI connectivity patterns predicted individual reaction times on corresponding face and scene perception tasks. Our findings demonstrate that the FFA and PPA anchor distinct intrinsic networks with unique spatio-temporal profiles that provide a functional architecture supporting their specialized roles in face and scene perception.

## Introduction

A fundamental principle of cortical organization is the existence of specialized networks dedicated to distinct cognitive domains (Fox & Raichle, 2007; Passingham et al., 2002; Power et al., 2011; Thomas Yeo et al., 2011). The processing of faces and scenes represents a canonical example of this functional specialization, supported by distinct yet interconnected neural systems in the human high-level visual cortex (Dilks et al., 2022; Epstein & Baker, 2019; Grill-Spector & Weiner, 2014; Haxby et al., 2000). Specifically, the Fusiform Face Area (FFA) and the Parahippocampal Place Area (PPA) are prominent regions that respond robustly to face and scene stimuli, respectively. The FFA, located in the fusiform gyrus, is critical for face recognition (Kanwisher & Yovel, 2006; Kanwisher et al., 1997; Parvizi et al., 2012; Puce et al., 1996; Schalk et al., 2017; Weiner & Zilles, 2016), and neuropsychological evidence from patients with prosopagnosia further emphasizes the functional specificity of the FFA (Barton et al., 2002; Damasio et al., 1982; McNeil & Warrington, 1993). Similarly, studies in macaques have highlighted the role of face-selective neurons in the homologous macaque FFA, which exhibit nearly exclusive responses to face stimuli (Tsao et al., 2006). The PPA, situated at the junction of the collateral and anterior lingual sulcus, plays a key role in scene perception and categorization (Dilks et al., 2022; Epstein & Baker, 2019; Epstein & Kanwisher, 1998), and serves broader functions, such as spatial navigation and contextual associations (Aminoff et al., 2013; Bar & Aminoff, 2003; Epstein, 2008). This functional specialization within the high-level visual cortex highlights the modular organization of face and scene processing, with the FFA specialized for face recognition and the PPA central to scene-related tasks.

While early research focused on identifying individual brain regions involved in face and scene perception using functional localizers (Epstein & Kanwisher, 1998; Kanwisher et al., 1997), subsequent efforts have shifted towards understanding the connectivity and interactions within distributed neural networks (Baldassano et al., 2016; Brandman & Peelen, 2017; Deen et al., 2024; Haxby et al., 2000; Peelen et al., 2009; Pitcher et al., 2011; Rajimehr et al., 2024; Singer et al., 2025; Watson & Andrews, 2024). For face perception, Haxby et al. (2000) proposed a foundational distributed model distinguishing between a ‘core system’ for visual analysis and an ‘extended system’ for further processing. The core system is comprised of the inferior occipital gyrus for early perception of facial features, the fusiform gyrus (FFA) for representing the invariant aspects of faces that underlie identity, and the superior temporal sulcus (STS) for representing changeable aspects like eye gaze and expression that facilitate social communication. Extended systems then process this information for functions like emotion recognition and directing spatial attention (Haxby et al., 2000). Similarly, scene perception relies on a distributed cortical network composed of three core regions: the PPA, the occipital place area (OPA), and the retrosplenial complex (RSC) (Epstein & Baker, 2019). This network is further organized into two distinct subnetworks (Baldassano et al., 2016; Epstein & Baker, 2019). A posterior network, including the OPA and posterior PPA, is strongly connected with visual and dorsal attention networks and is thought to process immediate visual features like spatial layout. In contrast, an anterior network, including the anterior PPA and RSC, connects robustly with the hippocampus, default mode, and frontoparietal control networks to support higher-level memory and navigational functions (Baldassano et al., 2016; Watson & Andrews, 2024). This functional and structural division is believed to support distinct computational goals such that the OPA supports visually guided navigation through the immediate environment, the RSC supports map-based navigation to out-of-sight locations, and the PPA, in a departure from earlier views, primarily supports scene categorization rather than navigation (Dilks et al., 2022).

Recent research indicates that the brain’s intrinsic functional and anatomical connectivity patterns can predict functional activations and relate to behavioral performance (Bedini et al., 2025; Gomez et al., 2015; Mantegna et al., 2025; Molloy et al., 2024; Saygin et al., 2012; Soyuhos & Baldauf, 2023; Zhu et al., 2011). For instance, Saygin et al. (2012) demonstrated that structural connectivity profiles of the fusiform gyrus could predict individual face-selective responses in the FFA, highlighting a strong link between anatomical connections and functional specialization. Building on this finding, Gomez et al. (2015) provided further evidence by identifying separate white-matter pathways associated with face- and place-selective regions and showing that local diffusion properties of these tracts correlated with category-specific behavioral performance. In addition to these anatomical findings, intrinsic functional connectivity patterns have proven highly informative. Zhu et al. (2011) found that resting-state functional connectivity strength between the occipital face area and the FFA correlated with individual differences in face recognition abilities, suggesting that intrinsic connectivity contributes to perceptual proficiency. Similarly, our recent study showed that mental imagery of faces versus scenes can be decoded from category-specific endogenously driven functional connectivity patterns (Mantegna et al., 2025). Together, these studies suggest that the brain’s intrinsic architecture, its structural and functional connectivity patterns, are linked to cognitive function and behavior. Understanding how specialized regions like the FFA and PPA are organized within these intrinsic networks is therefore important, as this underlying architecture is thought to provide the scaffold for efficient task-related processing and may explain individual differences in perceptual abilities (Fox & Raichle, 2007; Fox et al., 2005; Mennes et al., 2010; Zou et al., 2013).

In our study, we investigated the resting-state functional connectivity patterns of the FFA and PPA. Utilizing data from the Human Connectome Project (HCP), we employed both functional magnetic resonance imaging (fMRI) and magnetoencephalography (MEG) to characterize their intrinsic networks. First, using fMRI, we found that the FFA and PPA anchor spatially segregated intrinsic networks whose functional architecture aligns with their known task-based networks. Second, our MEG analyses revealed that this spatial segregation is reflected in the amplitude coupling of beta and gamma bands and uncovered frequency-specific directional dynamics, with incoming influences to these core regions carried by higher frequencies and outgoing influences by lower frequencies. Finally, we demonstrated that the strength of this intrinsic architecture is predictive of individual differences in behavior. Specifically, we established a functional double dissociation: connectivity patterns within the FFA-network predicted individual performance on a face-matching task, whereas connectivity within the PPA-network predicted performance on scene-processing tasks, but not vice-versa.

## Results

### Identification of FFA and PPA seed regions

We first identified the FFA and PPA seed regions within the multi-modal parcellation atlas (HCP-MMP1; (Glasser et al., 2016)) using a Neurosynth meta-analysis combined with a clustering approach (Figure 1A and Table S1, see Methods). For the ‘face’ association map, the largest cluster showed peak activation (MNI: +40.0, -50.0, -20.0) located within the FFC parcel, with its center of mass (MNI: +42.3, -59.7, -13.3) situated in the PH parcel, both in the right hemisphere. The second-largest ‘face’ cluster was localized in the left hemisphere, with both its peak activation (MNI: -40.0, -52.0, -22.0) and center of mass (MNI: -37.8, -65.5, -13.7) positioned within the FFC parcel. For the ‘place’ association map, the largest cluster, with both peak activation (MNI: -28.0, -46.0, -10.0) and center of mass (MNI: -29.7, -45.6, -10.3), was localized within the PHA3 parcel in the left hemisphere. The second-largest ‘place’ cluster exhibited peak activation (MNI: +28.0, -44.0, -14.0) within the VVC parcel, while its center of mass (MNI: +29.7, -43.0, -11.7) was localized to the PHA3 parcel in the right hemisphere (Table S1). These findings identified the FFC parcel as most frequently associated with face-related activations, and the PHA3 parcel as most frequently associated with scene-related activations, leading to their selection as seed regions for the FFA and PPA, respectively.

**Figure 1.**
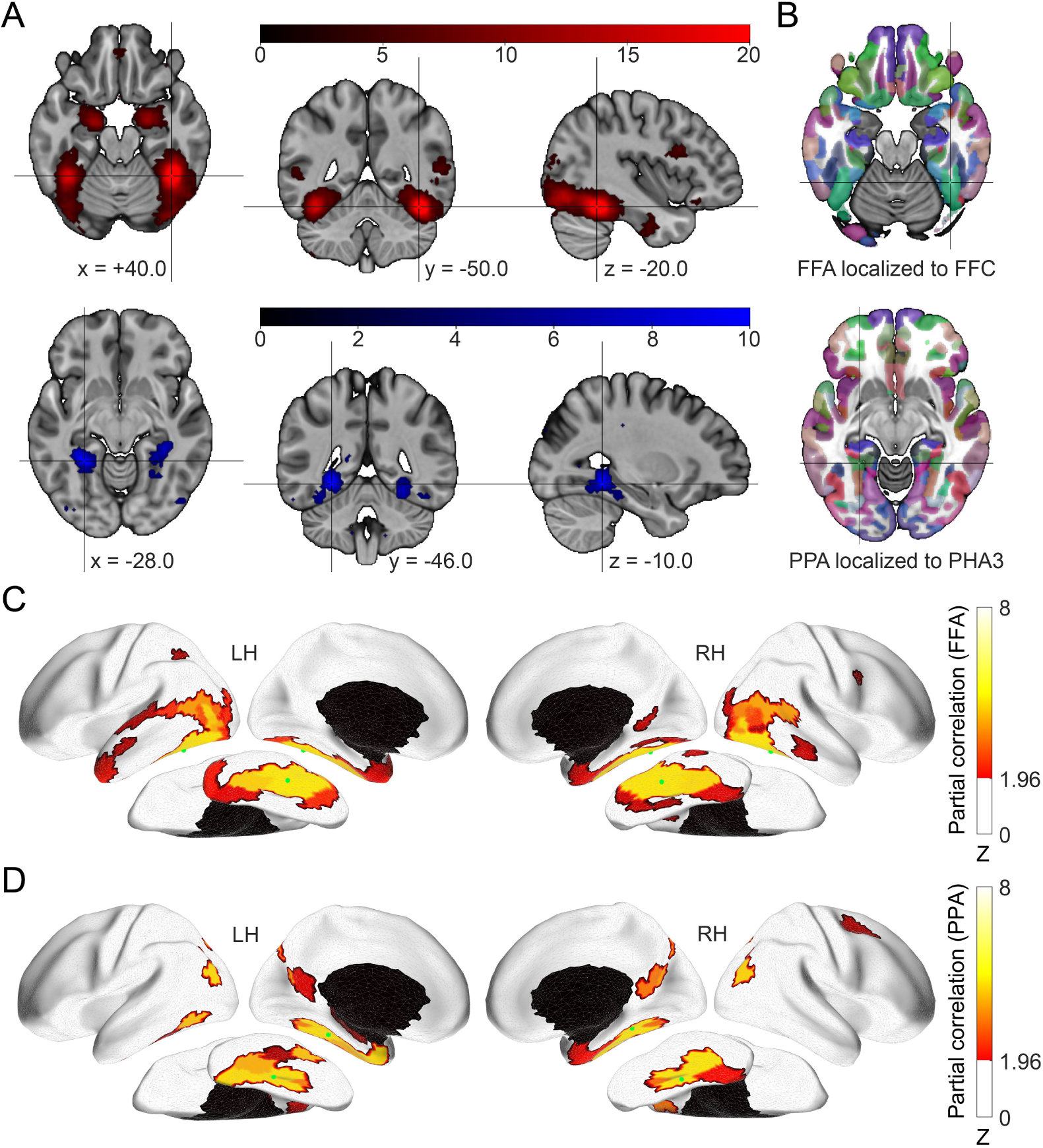
Anatomical locations and fMRI connectivity maps of the Fusiform Face Area (FFA) and Parahippocampal Place Area (PPA). (A) Neurosynth meta-analytic association maps generated for ‘face’ (top row, red) and ‘place’ (bottom row, blue) queries. For the ‘face’ map, the highlighted peak localizes the FFA to the FFC parcel in the right hemisphere (MNI coordinate in LPI order: +40.0, -50.0, -20.0). For the ‘place’ map, the highlighted peak localizes the PPA to the PHA3 parcel in the left hemisphere (MNI coordinate in LPI order: -28.0, -46.0, -10.0). (B) The location of the parcels in the Human Connectome Project Multi-Modal Parcellation (HCP-MMP1) atlas corresponding to the FFA and PPA. (C) Whole-brain resting-state fMRI functional connectivity map of the FFA seed region. (D) Whole-brain resting-state fMRI functional connectivity map of the PPA seed region. For panels (B) and (C), connectivity is displayed for the left (LH) and right (RH) hemispheres on each ipsilateral side. The green dots mark the seed regions. Statistical significance was assessed using a p < 0.05 threshold, FDR-corrected for 180 ROIs, and represented as z-scores.

### fMRI connectivity reveals distinct networks for FFA and PPA

Next, we examined the whole-brain fMRI connectivity patterns associated with these seeds. The FFA seed exhibited functional connectivity primarily with lateral occipital, ventral temporal, and lateral temporal cortical regions within each hemisphere (Figure 1C; see Table S2 for full list of significant connections). In contrast, the PPA seed showed strong connectivity largely with medial temporal lobe, posterior cingulate cortex, and inferior parietal cortex (Figure 1D; see Table S3 for full list). To identify regions selectively associated with face versus scene processing networks, we directly compared the connectivity profiles of the FFA and PPA seeds in each hemisphere (Wilcoxon signed-rank test, p < 0.05, FDR-corrected q < 0.05). This contrast revealed distinct sets of regions preferentially connected to either the FFA or the PPA (Figure 2). Specifically, 20 parcels demonstrated significantly stronger functional connectivity with the FFA seed region (Table 1). These FFA-preferential regions included areas within the visual cortex (V3B, V4), middle temporal complex (MT complex: MT, MST, PH, V4t), lateral occipital cortex (LO1-3), lateral temporal cortex (TE2p), STS (STSdp), temporoparietal junction (TPOJ1-3), ventral visual stream (PIT, V8, VVC), inferior parietal cortex (PFm), and premotor eye fields (PEF). Conversely, 13 parcels showed significantly stronger functional connectivity with the PPA seed region (Table 1). These PPA-preferential regions included areas within the ventral medial visual cortex (VMV2-3), adjacent parahippocampal cortex (PHA2), the entorhinal-perirhinal cortex (PeEc), medial and inferior parietal cortex (MIP, PGp, IP0), anterior and posterior cingulate cortex (25, DVT, POS1), superior premotor cortex (6a), and inferior frontal cortex (IFJp, IFSa). The specific parcels, their anatomical labels, and the statistics indicating the strength and significance of their preferential connectivity with either the FFA or PPA seed in each hemisphere are detailed in Table 1. These 33 differentially connected regions (20 FFA-preferential, 13 PPA-preferential) were subsequently used as target regions of interest (ROIs) for the MEG connectivity analyses.

**Figure 2.**
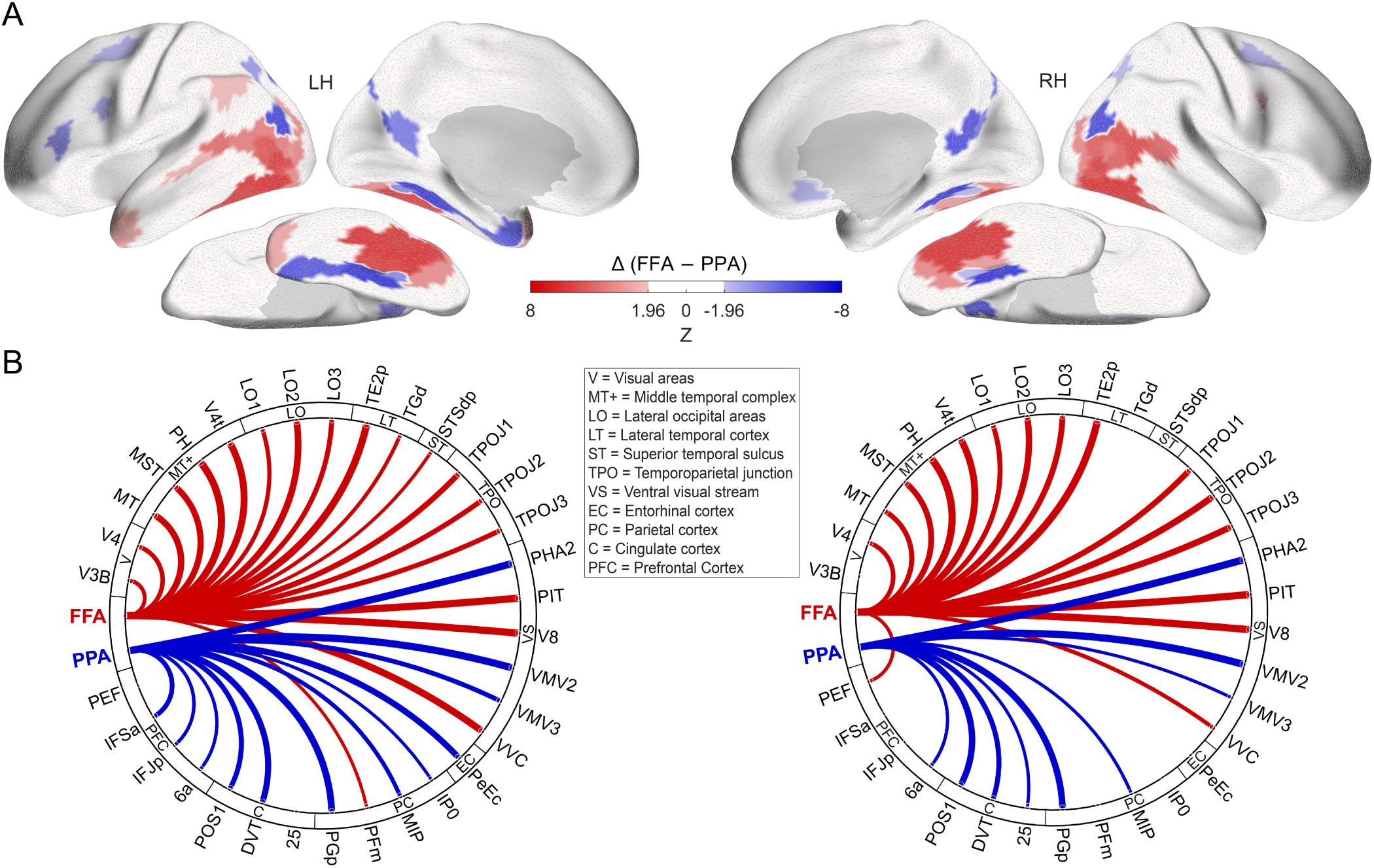
Contrasting fMRI connectivity profiles of the Fusiform Face Area (FFA) and Parahippocampal Place Area (PPA). (A) Surface maps display the direct contrast between the whole-brain connectivity profiles of FFA and PPA, with red indicating regions showing significantly stronger connectivity with FFA than PPA, and blue indicating regions with stronger connectivity with PPA than FFA. (B) Circular graphs for the left (LH) and right (RH) hemispheres illustrate the predominant functional connections between the FFA and PPA seed regions and the target regions from panel A. Red lines denote preferential connectivity with FFA, and blue lines with PPA; line thickness corresponds to the strength of this preferential connectivity (z-score). Statistical significance for these contrasts was assessed using a p < 0.05 threshold, FDR-corrected for 180 ROIs, with differences represented as z-scores. Abbreviations for the ROI groupings shown in the figure include: V = Visual areas, MT+ = Middle temporal complex, LO = Lateral occipital areas, LT = Lateral temporal cortex, ST = Superior temporal sulcus, TPO = Temporoparietal junction, VS = Ventral visual stream, EC = Entorhinal-perirhinal cortex, PC = Parietal cortex, C = Cingulate cortex, and PFC = Prefrontal Cortex.

**Table 1.**
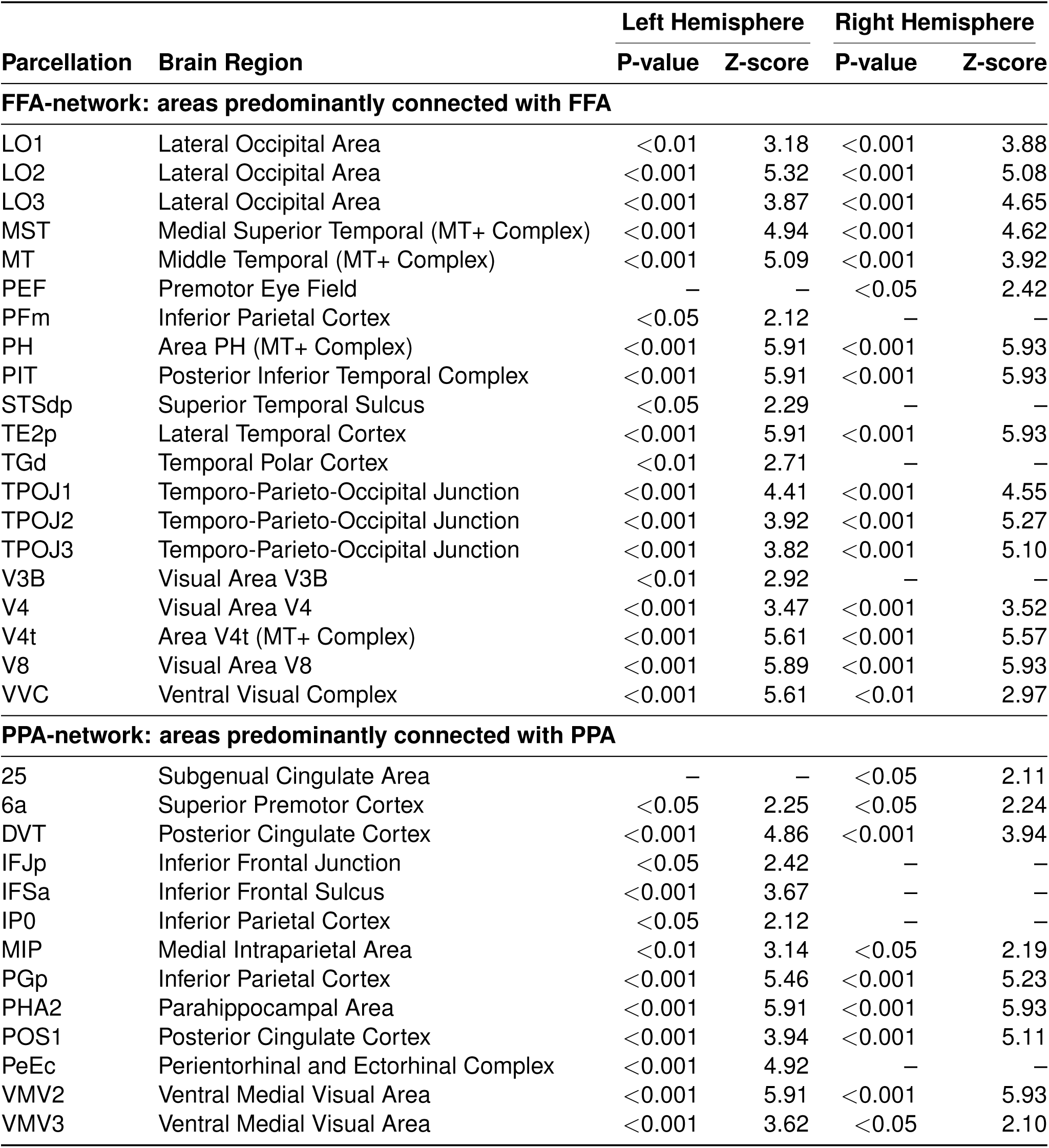
Brain regions exhibiting predominant functional connectivity with the Fusiform Face Area (FFA) and Parahippocampal Place Area (PPA). This table lists cortical regions from the Human Connectome Project multi-modal parcellation (HCP-MMP1) atlas, classifying each region according to its preferential connectivity with FFA or PPA (Figure 2). The ‘Parcellation’ column names the region, while the ‘Brain Region’ column provides an approximate functional label based on Glasser et al. (2013). Separate columns display the statistical significance (P-value) and strength (Z-score) of the functional connectivity for the left (LH) and right (RH) hemispheres. Dashes (–) indicate that significant connectivity was not detected in the corresponding hemisphere.

| Parcellation | Brain Region | Left Hemisphere |  | Right Hemisphere |  |
| --- | --- | --- | --- | --- | --- |
|  |  | P-value | Z-score | P-value | Z-score |
| FFA-network: areas predominantly connected with FFA |  |  |  |  |  |
| LO1 | Lateral Occipital Area | <0.01 | 3.18 | <0.001 | 3.88 |
| LO2 | Lateral Occipital Area | <0.001 | 5.32 | <0.001 | 5.08 |
| LO3 | Lateral Occipital Area | <0.001 | 3.87 | <0.001 | 4.65 |
| MST | Medial Superior Temporal (MT+ Complex) | <0.001 | 4.94 | <0.001 | 4.62 |
| MT | Middle Temporal (MT+ Complex) | <0.001 | 5.09 | <0.001 | 3.92 |
| PEF | Premotor Eye Field | – | – | <0.05 | 2.42 |
| PFm | Inferior Parietal Cortex | <0.05 | 2.12 | – | – |
| PH | Area PH (MT+ Complex) | <0.001 | 5.91 | <0.001 | 5.93 |
| PIT | Posterior Inferior Temporal Complex | <0.001 | 5.91 | <0.001 | 5.93 |
| STSdp | Superior Temporal Sulcus | <0.05 | 2.29 | – | – |
| TE2p | Lateral Temporal Cortex | <0.001 | 5.91 | <0.001 | 5.93 |
| TGd | Temporal Polar Cortex | <0.01 | 2.71 | – | – |
| TPOJ1 | Temporo-Parieto-Occipital Junction | <0.001 | 4.41 | <0.001 | 4.55 |
| TPOJ2 | Temporo-Parieto-Occipital Junction | <0.001 | 3.92 | <0.001 | 5.27 |
| TPOJ3 | Temporo-Parieto-Occipital Junction | <0.001 | 3.82 | <0.001 | 5.10 |
| V3B | Visual Area V3B | <0.01 | 2.92 | – | – |
| V4 | Visual Area V4 | <0.001 | 3.47 | <0.001 | 3.52 |
| V4t | Area V4t (MT+ Complex) | <0.001 | 5.61 | <0.001 | 5.57 |
| V8 | Visual Area V8 | <0.001 | 5.89 | <0.001 | 5.93 |
| VVC | Ventral Visual Complex | <0.001 | 5.61 | <0.01 | 2.97 |
| PPA-network: areas predominantly connected with PPA |  |  |  |  |  |
| 25 | Subgenual Cingulate Area | – | – | <0.05 | 2.11 |
| 6a | Superior Premotor Cortex | <0.05 | 2.25 | <0.05 | 2.24 |
| DVT | Posterior Cingulate Cortex | <0.001 | 4.86 | <0.001 | 3.94 |
| IFJp | Inferior Frontal Junction | <0.05 | 2.42 | – | – |
| IFSa | Inferior Frontal Sulcus | <0.001 | 3.67 | – | – |
| IP0 | Inferior Parietal Cortex | <0.05 | 2.12 | – | – |
| MIP | Medial Intraparietal Area | <0.01 | 3.14 | <0.05 | 2.19 |
| PGp | Inferior Parietal Cortex | <0.001 | 5.46 | <0.001 | 5.23 |
| PHA2 | Parahippocampal Area | <0.001 | 5.91 | <0.001 | 5.93 |
| POS1 | Posterior Cingulate Cortex | <0.001 | 3.94 | <0.001 | 5.11 |
| PeEc | Perientorhinal and Ectorhinal Complex | <0.001 | 4.92 | – | – |
| VMV2 | Ventral Medial Visual Area | <0.001 | 5.91 | <0.001 | 5.93 |
| VMV3 | Ventral Medial Visual Area | <0.001 | 3.62 | <0.05 | 2.10 |

### Frequency-specific MEG connectivity profiles of FFA and PPA

To investigate the temporal dynamics and directional interactions within the identified face- and place-preferential networks, we analyzed the MEG data focusing on connectivity between the FFA/PPA seed regions and the 33 target ROIs identified previously via fMRI (Table 1 and Figure 3A). We assessed amplitude and phase coupling using the orthogonalized power envelope correlation and the imaginary part of coherency metrics, respectively (Wilcoxon signed-rank test, p < 0.05, FDR-corrected q < 0.05).

**Figure 3.**
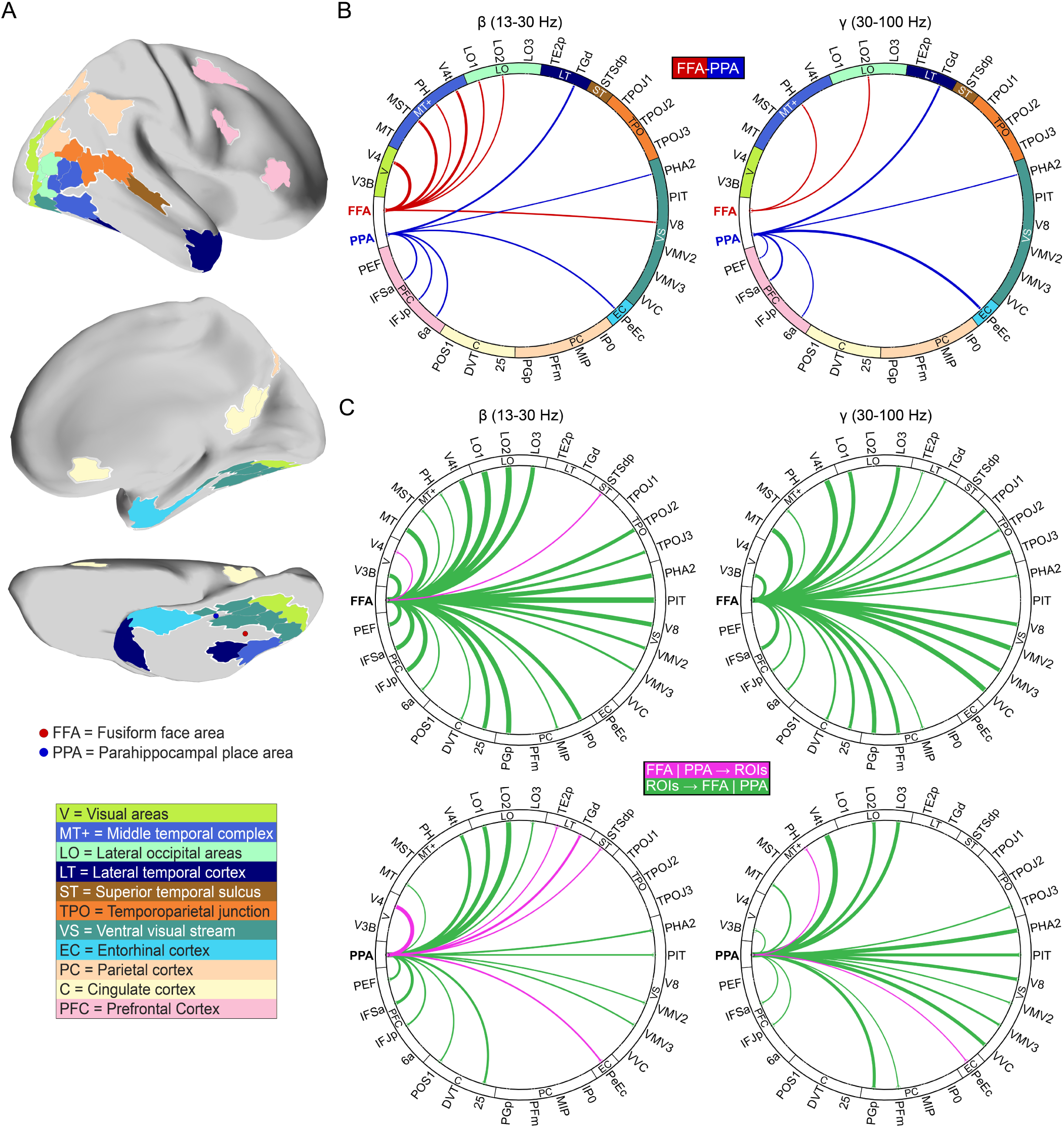
Frequency-specific connectivity profiles of the Fusiform Face Area (FFA) and Parahippocampal Place Area (PPA). (A) Location of regions of interest (ROIs) from Table 1 on cortex. The parcels with the same color have similar functional labels based on Glasser et al. (2013). (B) Predominant magnetoencephalography (MEG) functional connectivity profiles of the FFA and PPA to the areas shown in panel (A) in the beta (left) and gamma (right) frequency bands across two hemispheres. The results are based on the orthogonalized power envelope correlation method. Red and blue lines indicate preferential connectivity with the FFA and PPA, respectively. There were no significant regions in the other frequency bands. (C) Circular graphs show the directional connectivity in the beta and gamma bands for FFA (top) and PPA (bottom) with the regions in panel (A) across both hemispheres. For both graphs, magenta lines indicate significant outgoing influences (Seed → ROI), while green lines indicate significant incoming influences (ROI → Seed). The results for other frequency bands are in Figure S3. Statistical significance was assessed using a p < 0.05 threshold, FDR-corrected for 33 ROIs, and represented as z-scores. The width of the lines reflects the z-scores for each functional coupling. Abbreviations for the ROI groupings shown in the figure include: V = Visual areas, MT+ = Middle temporal complex, LO = Lateral occipital areas, LT = Lateral temporal cortex, ST = Superior temporal sulcus, TPO = Temporoparietal junction, VS = Ventral visual stream, EC = Entorhinal-perirhinal cortex, PC = Parietal cortex, C = Cingulate cortex, and PFC = Prefrontal Cortex.

Analysis of amplitude coupling in the beta (13-30 Hz) and gamma (30-100 Hz) frequency bands reflected the distinct network segregation observed with fMRI (Figure 3B). In both frequency bands, the FFA seed showed significantly stronger connectivity with target ROIs located primarily in the lateral occipital cortex, MT complex, and earlier visual areas. Conversely, the PPA seed exhibited significantly stronger connectivity with target ROIs in the entorhinal-perirhinal cortex and the posterior prefrontal cortex in both beta and gamma bands (see Figure S1 for whole-brain connectivity maps). However, such distinct network segregation with amplitude coupling was not evident in the lower frequency bands; neither the FFA nor PPA seeds demonstrated significant preferential connectivity to these target ROIs in the delta (1–4 Hz), theta (4–8 Hz), or alpha (8–13 Hz) bands. In contrast, phase-based connectivity analyses only revealed significant preferential connectivity in the beta (13-30 Hz) band, specifically between the PPA and the posterior inferior temporal cortex (PIT) (see Figure S2 for whole-brain connectivity maps).

We next examined frequency-specific directional interactions using the partial directed coherence (PDC) method across two hemispheres (Wilcoxon signed-rank test, p < 0.05, FDR-corrected q < 0.05) (Figure 3C and S3). Incoming influences (ROI → Seed), directed towards the FFA and PPA seeds, were predominant in the higher frequencies (beta and gamma) (green lines, Figure 3C). Prominent sources driving these high-frequency incoming influences included the lateral occipital cortex (LO1-3), MT complex (MT, V4t), ventral visual stream areas (PHA2, PIT, V8, VMV2-3, VVC), temporoparietal junction (TPOJ: TPOJ2-3), and PFC regions (PEF, IFSa, IFJp, 6a). Notably, while contributing significantly to these highfrequency interactions, influences originating from PFC specifically exhibited their peak strength in the low frequency band. Conversely, outgoing influences (Seed → ROI), originating from the FFA and PPA seeds, were relatively stronger in the lower frequencies (delta, theta, and alpha) (magenta lines, Figure S3). Significant outgoing influences were directed towards target ROIs including area V4, STS (STSdp), TPJ (TPOJ1), ventral visual cortex (VVC), the entorhinal-perirhinal cortex (PeEc), and posterior cingulate cortex (DVT, POS1). Overall, incoming influences were most robust in the gamma band, while outgoing influences were most robust in the alpha band.

In summary, MEG analyses further characterized the spatial segregation of FFA and PPA networks observed in fMRI, particularly within beta and gamma band amplitude coupling. Additionally, these analyses revealed frequency-specific temporal dynamics characterized by predominant incoming influences in high frequency bands (beta, gamma) and relatively stronger outgoing influences in low frequency bands (delta, theta, and alpha).

### Intrinsic network connectivity predicts behavioral performance

Finally, we investigated whether the distinct intrinsic functional connectivity patterns of the FFA and PPA networks were related to individual differences in behavioral performance on relevant tasks. We employed the connectome-based predictive modeling (CPM) using fMRI partial correlations within the previously defined FFA-preferential network (37 regions including bilateral seeds and significant ROIs from Table 1) and PPA-preferential network (23 regions including bilateral seeds and significant ROIs from Table 1). These networks were used to predict median reaction times (RTs) from the face-matching task and the 0-back and 2-back scene tasks, respectively (task details in Figure 4). All models were rigorously tested using leave-one-out cross-validation and permutation testing (1000 iterations) for statistical significance.

**Figure 4.**
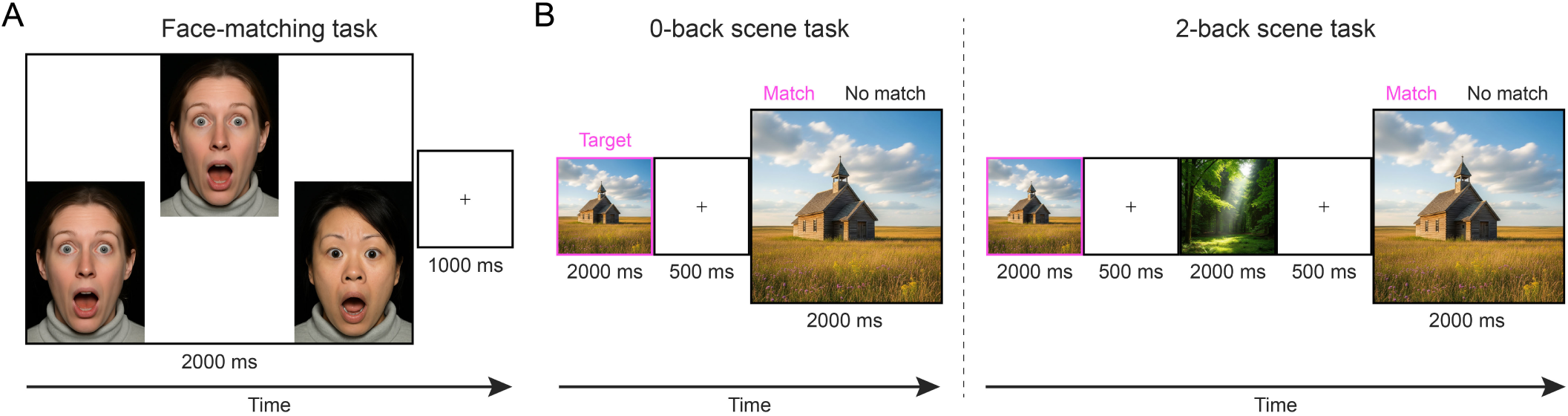
Face-matching and n-back scene paradigms. (A) In the face-matching task, participants identified which of two choice faces matched a target face. Each trial consisted of a 2000 ms stimulus presentation followed by a 1000 ms fixation interval. (B) In the scene n-back task, participants performed either a 0-back (left) or 2-back (right) working memory task. For both conditions, each stimulus was presented for 2000 ms followed by a 500 ms fixation interval. The illustrative images shown here were computer generated.

First, we tested if connectivity within the FFA network could predict performance on the face-matching task (n = 350 participants). The CPM analysis revealed a significant positive correlation between predicted and actual median RTs (Spearman’s r = 0.31, p = 0.014, permutation test; Figure 5A), indicating that stronger intrinsic connectivity patterns within this network were associated with faster face-matching performance. In contrast, a control model using connectivity within the PPA network failed to predict face-matching RTs (Spearman’s r = -0.09, p = 0.707, permutation test; Figure 5B), demonstrating the specificity of the FFA network’s relationship with face processing behavior. The successful prediction was driven by a distributed set of connections consistently selected as predictive features across subjects (Figure 5C), with regions within the MT complex (MST, MT, PH, V4t) and inferior parietal cortex (PFm) showing the highest number of predictive pairs within the FFA network.

**Figure 5.**
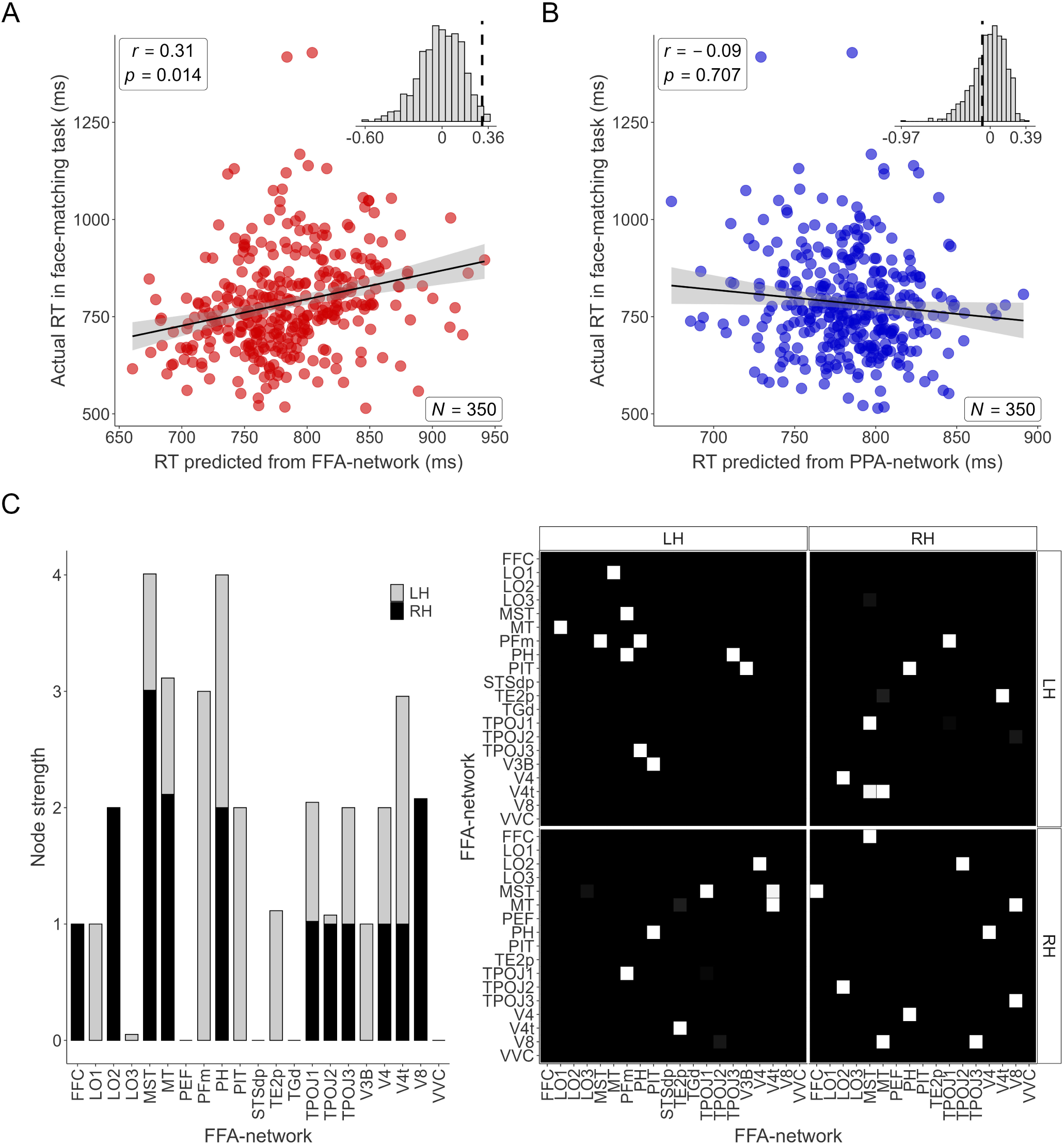
Connectome-based predictive modeling (CPM) of median reaction times in the face-matching task. (A) The scatter plot compares actual median reaction times (RT; ms) with those predicted by CPM, based on resting-state fMRI connectivity within the FFA-network (37 regions, including bilateral seeds and significant ROIs from Table 1). The black line represents the linear regression fit, and the gray shading indicates the 95% confidence interval. The inset histogram displays the null distribution of correlations obtained through permutation testing, with the dashed vertical line marking the observed correlation. (B) The same procedure as in (A) but using a CPM model based on connectivity within the PPA-network (23 regions, including bilateral seeds and significant ROIs from Table 1). (C) The connectivity pairs driving the successful prediction shown in (A). The left bar chart displays node strength, a measure of each region’s total contribution to the predictive model. Node strength for a given region is calculated by summing the selection consistency scores (the values from the matrix on the right) for all connections involving that region. The adjacency matrix on the right visualizes these connections; the brightness of each cell represents the proportion of cross-validation folds in which that edge was selected as a significant predictor of reaction time.

Next, we assessed the relationship between PPA network connectivity and performance on the scene n-back tasks (n = 352 participants). Connectivity within the PPA network significantly predicted median RTs for the 0-back scene task (Spearman’s r = 0.25, p = 0.036, permutation test; Figure 6A). The control model using the FFA network connectivity did not significantly predict 0-back RTs (Spearman’s r = - 0.02, p = 0.546, permutation test; Figure 6E), further supporting network specificity. Similarly, connectivity within the PPA network also significantly predicted median RTs for the more demanding 2-back scene task (Spearman’s r = 0.29, p = 0.011, permutation test; Figure 6C). The corresponding control model using FFA network connectivity failed to predict 2-back RTs (Spearman’s r = 0.05, p = 0.388, permutation test; Figure 6F). For both the 0-back and 2-back scene tasks, the successful predictions using the PPA network were based on distributed patterns of connectivity (Figure 6B, 6D). Notably, the specific patterns differed: the 0-back prediction drew heavily on connections within parahippocampal (PHA2, PHA3), posterior cingulate (POS1, DVT), and ventral medial visual (VMV3) cortex (Figure 6B), while the 2-back prediction showed increased reliance on connections involving frontal (6a, IFJp) and parietal (PGp, MIP) regions (Figure 6D). Taken together, these CPM results demonstrate a double dissociation: intrinsic resting-state functional connectivity within the FFA-network specifically predicted individual differences in face-matching performance, whereas connectivity within the PPA-network specifically predicted performance on both 0-back and 2-back scene tasks, highlighting a significant link between the brain’s intrinsic network architecture and domain-specific perceptual processing capabilities.

**Figure 6.**
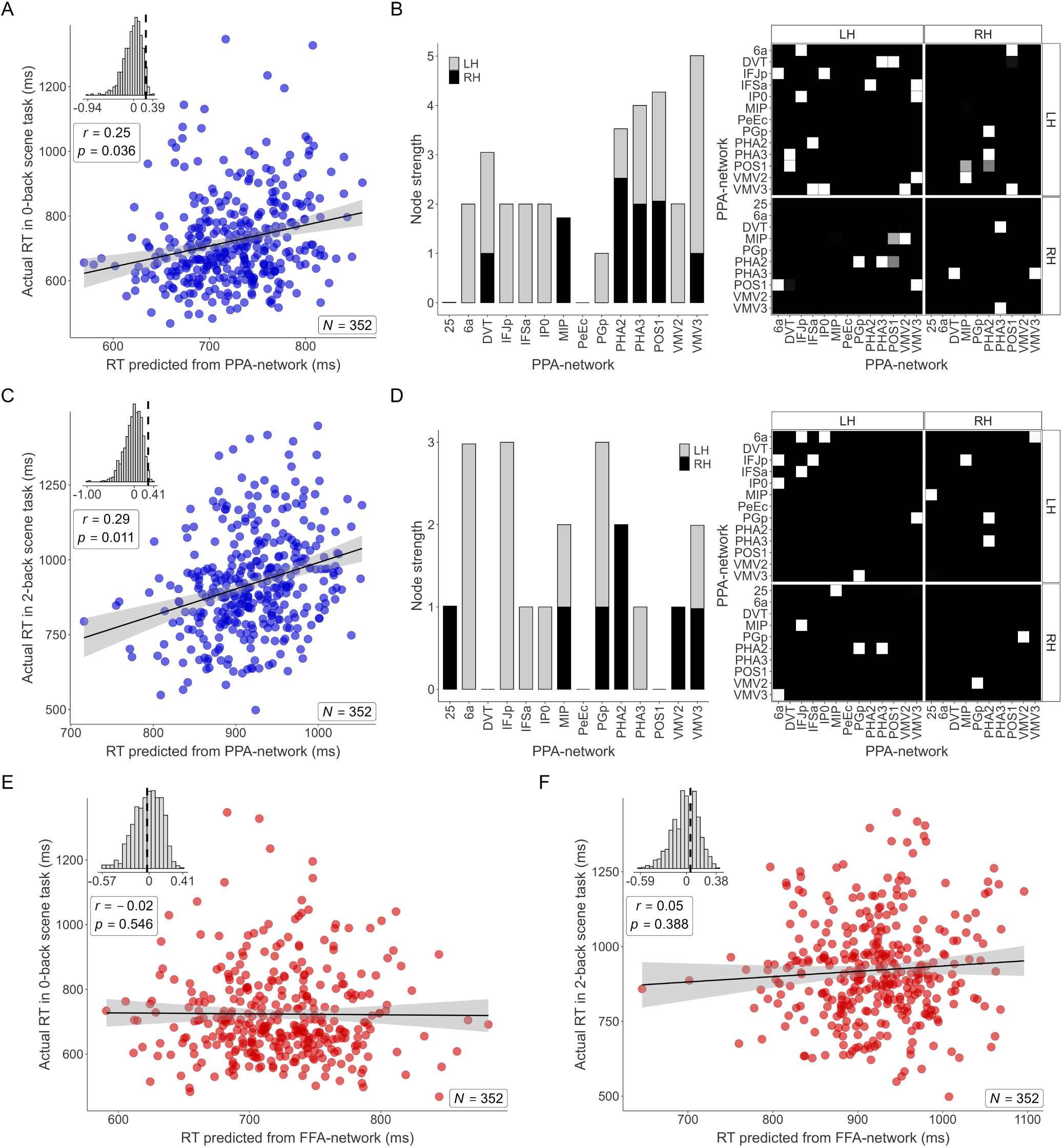
Connectome-based predictive modeling (CPM) of median reaction times in 0-back and 2-back scene tasks. (A) The scatter plot compares actual median reaction times (RTs; ms) from 0-back scene task with those predicted by CPM, based on resting-state fMRI connectivity within the PPA network (23 regions, including bilateral seeds and significant ROIs from Table 1). The black line represents the linear regression fit, and the gray shading indicates the 95% confidence interval. The inset histogram displays the null distribution of correlations obtained through permutation testing, with the dashed vertical line marking the observed correlation. (B) The connectivity pairs driving the successful prediction shown in (A). The left bar chart dis- plays node strength, a measure of each region’s total contribution to the predictive model. Node strength for a given region is calculated by summing the selection consistency scores (the values from the matrix on the right) for all connections involving that region. The adjacency matrix on the right visualizes these connections; the brightness of each cell represents the proportion of cross-validation folds in which that edge was selected as a significant predictor of reaction time. (C) Scatter plot and (D) connectivity pairs for predicting the 2-back scene task RTs using the same PPA network as in (A). (E) The same procedure as in (A) but predicting the 0-back scene task RTs using a CPM model based on connectivity within the FFA network (37 regions, including bilateral seeds and significant ROIs from Table 1). (F) Similar to (E), but predicting the 2-back scene task RTs using the FFA network.

## Discussion

This study provides compelling evidence that the core regions for face (FFA) and scene (PPA) perception are embedded within distinct intrinsic functional networks. These networks are characterized by unique spatial and temporal connectivity profiles that are present even during task-free rest. Leveraging the complementary strengths of fMRI and MEG, we demonstrated three core findings: First, fMRI revealed spatially segregated networks primarily characterized by the FFA’s preferential connectivity with lateral occipitotemporal, inferior temporal, and temporoparietal regions versus the PPA’s connectivity with ventral medial visual, posterior cingulate, and entorhinal-perirhinal areas. Second, MEG analyses showed that this spatial segregation is mirrored in the amplitude coupling of higher frequency bands (beta, gamma), and further uncovered frequency-specific directional dynamics, highlighting dominant incoming influences (ROI → Seed) in higher frequencies and relatively stronger outgoing influences (Seed → ROI) in lower frequencies. Third, our predictive model established a direct behavioral relevance for this intrinsic architecture: connectivity patterns within the FFA-network specifically predicted individual face-matching performance, whereas connectivity within the PPA-network specifically predicted performance on both 0- and 2-back scene tasks, and not vice versa, demonstrating a clear double dissociation.

The distinct fMRI connectivity profiles observed here strongly reinforce the principle that the functional specialization of the FFA and PPA (Epstein & Kanwisher, 1998; Kanwisher et al., 1997) is embedded within the brain’s intrinsic functional architecture, resonating with extensive work showing correspondence between resting-state and task-based networks (Cole et al., 2014; Fox & Raichle, 2007; Smith et al., 2009; Tavor et al., 2016) and confirming this intrinsic organization provides a scaffold for efficient perceptual processing. Specifically, the FFA’s preferential connectivity with the lateral occipital cortex, MT complex, STS, and TPJ aligns well with established core and extended face processing networks (de Vries & Baldauf, 2019; Duchaine & Yovel, 2015; Haxby et al., 2000; Mü ller et al., 2018; Rajimehr et al., 2009, 2024; Von Der Heide et al., 2013). Previous models proposed a division between the FFA for invariant identity processing and the posterior STS (pSTS) for changeable aspects like expression (de Vries & Baldauf, 2019; Gobbini & Haxby, 2007; Haxby et al., 2000), with extensions highlighting pSTS’s role in dynamic identity cues (O’Toole et al., 2002). However, a more recent perspective emphasizes a primary functional division between form processing via the ventral stream and motion processing via the dorsal stream (Bernstein & Yovel, 2015). Our connectivity results fit well within this evolving understanding: the FFA’s connection with the lateral occipital cortex underscores their role as a ventral pathway analyzing facial form; connectivity with the MT complex might reflect interaction with general motion analysis that interfaces with the specialized processing of facial motion (like expression or dynamic identity) in the STS regions; and the linkage to the TPJ could represent the integration of identity perception with the retrieval of person knowledge and theory of mind processes (Gobbini & Haxby, 2007). This revised view, emphasizing form versus motion, also accommodates findings that the FFA processes expression (likely its form components) and the pSTS responds to static cues implying motion (Bernstein & Yovel, 2015), highlighting the integrated nature of face perception supported by the observed intrinsic connectivity network.

Conversely, the PPA network’s preferential connectivity with adjacent parahippocampal cortex, inferior parietal areas, and posterior cingulate cortex map onto established networks for scene recognition, spatial navigation, and contextual memory integration (Aminoff et al., 2013; Bar & Aminoff, 2003; Caspers et al., 2013; Epstein, 2008; Epstein & Baker, 2019; Rajimehr et al., 2024). Of particular significance in our results is the strong coupling observed between the PPA and the entorhinal-perirhinal cortex. This connection likely reflects the critical interplay between scene-level processing (Epstein & Baker, 2019) and the detailed object processing functions attributed to the entorhinal-perirhinal cortex, especially the perirhinal cortex (Brown & Aggleton, 2001; Deshmukh et al., 2012; Murray & Bussey, 1999; Suzuki & Naya, 2014). The perirhinal cortex is crucial for representing object features, identifying objects (Murray & Bussey, 1999), and signaling item familiarity (Brown & Aggleton, 2001; Suzuki & Naya, 2014). Furthermore, The perirhinal cortex supports various forms of associative memory, linking items together or associating items with rewards or context (Murray & Bussey, 1999; Suzuki & Naya, 2014). Notably, the lateral entorhinal cortex receives input from the perirhinal cortex (Suzuki & Naya, 2014), and represents spatial information derived from landmarks when objects are present, suggesting an integration of object and spatial context (Deshmukh et al., 2012). Therefore, the strong functional connectivity between PPA and the entorhinal-perirhinal cortex observed here likely facilitates the integration of object identity and familiarity information (Brown & Aggleton, 2001; Deshmukh et al., 2012; Murray & Bussey, 1999; Suzuki & Naya, 2014) with the broader spatial layout and context represented by PPA (Epstein & Baker, 2019). This interaction might be essential for building holistic scene representations, enabling functions such as recognizing objects within their typical scenes, associating objects with specific locations (Brown & Aggleton, 2001), and potentially using familiar objects as landmarks (Epstein & Baker, 2019) within a scene context processed by PPA.

The MEG analyses provided further insights into the temporal and directional dynamics governing communication within these intrinsic networks. First, we found that amplitude coupling in the beta (13-30 Hz) and gamma (30-100 Hz) frequency bands mirrored some important aspects of the spatial segregation observed with fMRI. This reinforces the role of these faster oscillations in maintaining information processing within these specialized resting-state networks, potentially binding distributed neural populations (de Pasquale et al., 2010; Engel et al., 2001; Hipp et al., 2012; Siegel et al., 2012). Other frequency bands, such as alpha, theta and delta, did not show such a spatial congruence with the fMRI results. Second, the directionality analysis revealed frequency-specific directed interactions between the FFA/PPA seeds and their associated ROIs. Importantly, interpreting these directed influences requires considering the specific anatomical pathways involved, as FFA and PPA regions function as mid-level hubs in the functional hierarchy. For example, the signals arriving at these seeds (ROI → Seed) can represent feedforward input if originating from lower and parallel visual areas (e.g., V4t, LO1-3) or feedback if descending from higher cortical regions (e.g., from PFC, TPJ). Similarly, the signals departing from these seeds (Seed → ROI) can represent feedback (e.g., to V4) or feedforward projections (e.g., to STS, entorhinal-perirhinal cortex). Within this anatomical context, our findings indicated a general principle: incoming influences were predominantly carried by higher frequencies (beta and gamma), integrating signals from diverse cortical sources. In contrast, outgoing influences were relatively stronger in lower frequencies (delta, theta, and alpha), projecting towards both lower and higher processing stages. This frequency-specific directional architecture may form the intrinsic basis for the temporal dynamics observed during active perception. For instance, MEG decoding has shown that the brain begins to process face-specific information exceptionally early around 61 ms (Cichy et al., 2014). Notably, this process unfolds in a coarse-to-fine sequence; the brain extracts categorical information like gender starting around 72 ms, before recognizing a face’s specific identity at 91 ms (Dobs et al., 2019). In contrast, feedback-driven contextual processes, such as a scene facilitating the perception of an object, occur significantly later around 320 ms (Brandman & Peelen, 2017). Further investigation is needed to understand how the frequency-specific directed interactions we observed during rest map onto these task-evoked temporal dynamics.

Finally, we showed that this intrinsic functional architecture, measured during task-free rest, is directly and specifically linked to individual cognitive abilities. The CPM results provide robust, cross-validated evidence for this link, successfully predicting individual differences in face-matching and scene n-back RTs from fMRI connectivity patterns within the functionally relevant networks. Crucially, the observed double dissociation – the FFA network predicting only face task performance and the PPA network predicting only scene task performance – offers strong evidence that functional specificity is encoded within these intrinsic networks. This work significantly extends previous correlational studies (Zhu et al., 2011) by employing a rigorous predictive framework (Finn et al., 2015; Rosenberg & Finn, 2022; Shen et al., 2017) and demonstrating this specificity across both face and scene domains within the same participants. Moreover, the distinct connectivity patterns within the PPA network predicting performance of the 0-back versus the more demanding 2-back scene task suggest that subtle variations in intrinsic network organization are sensitive to, and predictive of, the capacity to engage network resources differently based on cognitive load. The increased reliance on connectivity involving frontal (6a, IFJp) and parietal (PGp, MIP) regions for predicting 2-back performance likely reflects the intrinsic substrate supporting the greater recruitment of working memory and executive control needed for this higher-load condition (Baldauf & Desimone, 2014; Mantegna et al., 2025). This frontoparietal involvement may also support the implementation of top-down search strategies that create a category-specific bias for task-relevant objects in visual cortex (Peelen et al., 2009), with parietal regions like the intraparietal sulcus thought to be a source of these contextual guidance signals (Peelen, 2025).

Importantly, our findings stem from resting-state data, and directly comparing these intrinsic patterns with connectivity dynamics during active face and scene perception using both fMRI and MEG is a crucial next step to fully understand task-dependent network modulation. Additionally, assessing the generalizability of these findings beyond the healthy young adult HCP sample requires investigating these network profiles across diverse ages and in clinical populations with known perceptual differences (e.g., prosopagnosia, autism spectrum disorder). Such research would allow exploration of potential interactions across domains (face/scene) and provide further tests of the functional specificity and dissociation of the underlying FFA and PPA networks. Addressing these points, alongside exploring more advanced analyses like dynamic functional connectivity or cross-frequency coupling, promises to significantly enhance our understanding of how these specialized networks dynamically support face and scene perception.

In conclusion, our results show that the FFA and PPA anchor distinct intrinsic functional networks, characterized not only by unique spatial topographies but also by specific temporal and directional dynamics observable in frequency-specific MEG signals. Critically, we established that this intrinsic network architecture, present during task-free rest, is functional as connectivity patterns within these networks specifically predict individual differences in face and scene processing abilities. These findings underscore the power of integrating multimodal neuroimaging to reveal the brain’s specialized functional organization and understand the fundamental link between the brain’s intrinsic spatio-temporal network structure and human cognitive function.

## Methods

### Participants and dataset

We used data from the HCP 1200 Subjects Release (Van Essen et al., 2013) for our analyses. This comprehensive dataset includes protocols involving resting-state functional MRI (fMRI), resting-state MEG, and structural MRI (Barch et al., 2013; Larson-Prior et al., 2013; Smith et al., 2013a). For the primary fMRI and MEG connectivity analyses, we initially considered 95 participants who completed the MEG protocol. From this group, a subset of 55 individuals (26 female, aged 22–35) was selected to avoid family-based confounds by including only one member from each twin pair, following procedures similar to Soyuhos and Baldauf (2023). For the subsequent CPM analyses, we used data from a larger cohort of 371 subjects (192 female, aged 22–36) selected from the HCP 1200 release, ensuring that each participant was from a different family structure. All participants included in these analyses were healthy adults with no reported neurological or psychiatric disorders, and informed consent was obtained prior to participation. Further details on inclusion and exclusion criteria for the HCP cohort can be found in Van Essen et al. (2013). The anonymized dataset is available via ConnectomeDB (Hodge et al., 2016; Marcus et al., 2013).

### Behavioral measures

Behavioral data were derived from tasks administered as part of the HCP task fMRI battery (Barch et al., 2013). For this study, we focused on behavioral scores from a face-matching task and a scene n-back task. In the face-matching task, participants viewed a target face presented above two alternative faces (Figure 4A). The faces displayed either an angry or fearful expression. Participants then indicated via button press which alternative (left or right) matched the target. The scene n-back task alternated between 0-back and 2-back conditions, signaled by on-screen prompts (Figure 4B). In the 0-back condition, participants indicated via button press whether the currently displayed scene matched the predefined target shown for that task. In the 2-back condition, participants indicated whether the current scene matched the one shown two images previously or not. For the CPM analysis, we used the median RT from correct trials as the behavioral score, calculated separately for the face-matching, 0-back scene, and 2-back scene tasks.

### Acquisition and preprocessing of resting-State fMRI and MEG data

The fMRI data were obtained from the minimally preprocessed CIFTI dense time series available on ConnectomeDB to ensure data reproducibility (Glasser et al., 2013; Smith et al., 2013a). Imaging was performed using a customized Siemens 3T Connectome Skyra scanner equipped with a 32-channel head coil and body transmission coil. T1-weighted structural images were collected using a 3D MPRAGE sequence with 0.7 mm isotropic resolution (TR = 2400 ms; TE = 2.14 ms; TI = 1000 ms; flip angle = 8°). The data consisted of four runs, each approximately 15 minutes long. During acquisition, participants were instructed to maintain fixation on a crosshair presented on a dark background. Imaging parameters for fMRI included: TR = 720 ms, TE = 33.1 ms, flip angle = 52°, field of view (FOV) = 208 × 180 mm, matrix size = 104 × 90, slice thickness = 2 mm, and a multi-band factor of 8. The HCP minimal preprocessing pipeline for fMRI involved correction for spatial distortions, head motion, and B0 field inhomogeneities. Functional images were registered to the T1-weighted structural image and subsequently normalized to 2 mm MNI space. Following normalization, the functional data were resampled to a standard mesh and integrated into a standard gray ordinates space using MSMAll registration. This approach ensured consistent alignment of cortical and subcortical regions across participants, facilitating accurate group-level analysis (Glasser et al., 2013; Smith et al., 2013a).

The MEG data were collected using a Magnes 3600 whole-head scanner featuring 248 magnetometers and 23 reference channels. Participants were recorded in a supine position while fixating on a red crosshair. Data were sampled at a rate of 2034.5101 Hz and stored in a 4D file format. Electrooculography, electrocardiography, and electromyography were recorded concurrently to aid in artifact correction. Preprocessing for MEG followed methods described by Soyuhos and Baldauf (2023). The Preprocessed Resting-State package from ConnectomeDB provided annotations identifying bad channels, bad segments, and non-brain artifact components (Hodge et al., 2016). Artifacts, such as those arising from eye blinks and muscle activity, were identified and corrected using functions within the FieldTrip toolbox (Oostenveld et al., 2011). The MEG data were then bandpass filtered between 1.3–150 Hz, and notch filters were applied to eliminate 60 Hz and 120 Hz power line noise.

### Resting-state fMRI and MEG connectivity analysis

To calculate functional connectivity from the fMRI data, we followed the steps outlined in Smith et al. (2013a). First, the minimally preprocessed CIFTI dense time series were normalized ((x - mean) / stdev). Data from all four runs for each participant were then concatenated into a single continuous dataset (one hour per subject). Brain regions were defined using the HCP-MMP1 atlas (Glasser et al., 2016). A ‘dense connectome’ matrix was subsequently computed using the Connectome Workbench toolbox, representing the partial correlations between all 360 regions in the atlas. The partial correlation approach was chosen because it aims to infer direct connections between node pairs by statistically removing the influence of time series from all other network nodes before calculating the correlation between the pair in question. This method thereby provides a more accurate estimate of the functional connectome compared to full correlation (Smith et al., 2013b). Seed-based exploratory and ROI-based connectivity analyses were then performed separately for the left and right seed regions within their respective hemispheres.

For the MEG data, frequency-specific functional connectivity matrices were calculated following the steps detailed in Soyuhos and Baldauf (2023). We began by reconstructing source-level activity using the Brainstorm toolbox (Tadel et al., 2011), mapping the activity onto each subject’s native cortical surface. Specifically, minimum-norm estimation (MNE) was used to estimate cortical activity at 15,002 distributed sources across the cortex (Baillet et al., 2001; Dale et al., 2000; Hä mä lä inen & Ilmoniemi, 1994). This source-level data was then parcellated into the same 360 regions defined by the HCP-MMP1 atlas (Glasser et al., 2016). To account for the distinct contributions of phase- and amplitude-coupling to functional connectivity (Daffertshofer et al., 2018; Siems & Siegel, 2020), we employed both phase- and amplitude-based connectivity measures. Specifically, we used the imaginary part of coherency (Nolte et al., 2004) to assess phase relationships and the orthogonalized power envelope correlation (Hipp et al., 2012) to measure amplitude-based connectivity. These metrics were selected for their effectiveness in reducing spatial leakage artifacts and enhancing the consistency of group-level analyses (Bastos & Schoffelen, 2016; Colclough et al., 2016; Duan et al., 2021).

Furthermore, to analyze dominant directional interactions between our seed regions (FFA, PPA) and target regions of interest (ROIs) during the resting state, we computed PDC (Baccalá & Sameshima, 2001). PDC values were first calculated for each unidirectional connection between seed regions and target ROIs. Then, for each pair, the dominant direction of influence was determined by statistically comparing the PDC values for the opposing connections (Seed → ROI vs. ROI → Seed) to identify the significantly stronger direction. PDC was selected based on several advantages: it primarily considers direct interactions, is generally regarded as relatively insensitive to source leakage, and exhibits high group-level repeatability compared to alternative directionality measures (Colclough et al., 2016). All MEG functional connectivity analyses were performed across distinct frequency bands, including delta (1–4 Hz), theta (4–8 Hz), alpha (8–13 Hz), beta (13–30 Hz), and gamma (30–100 Hz), to assess frequency-specific interactions between regions.

### Identifying seed and target ROIs

To determine the specific parcels in the HCP-MMP1 atlas corresponding to our seed regions for the FFA and PPA, we first generated meta-analytic association maps using Neurosynth (Yarkoni et al., 2011). These maps identified regions consistently activated across numerous studies: an analysis for the term ‘face’ included 896 studies (31842 activation foci), and an analysis for ‘place’ included 189 studies (6595 activation foci). These Neurosynth association maps were generated in MNI152 2mm space and represented statistically significant regions of consistent activation (FDR-corrected, q < 0.01). We then used the AFNI toolbox (Cox, 1996) to identify spatially contiguous clusters of these significant voxels using a nearest-neighbor algorithm where voxels were grouped if their faces touched. The top two largest by volume (in voxels) for each map (‘face’ and ‘place’) were extracted. These clusters’ MNI coordinates for both their peak activation (Peak; defined as the voxel with the highest statistical value within the cluster) and their center of mass (CMass; representing the average spatial location of the cluster) are presented in Table S1. MNI coordinates are reported in LPI order, where positive X, Y, and Z values correspond to anatomical Right, Anterior, and Superior, respectively. Subsequently, these extracted Peak and CMass MNI coordinates were localized by referencing them against the volumetric version of the HCP-MMP1 atlas (Bedini et al., 2023).

The target ROIs were selected using a data-driven approach designed to isolate areas exhibiting distinct functional connectivity profiles relative to the FFA and PPA seed regions. We performed statistical tests on subject-specific fMRI partial correlation connectivity maps. Specifically, for each parcel in the MMP1 atlas within each hemisphere, we compared its connectivity strength to the ipsilateral FFA seed region versus its connectivity strength to the ipsilateral PPA seed region across subjects. Parcels exhibiting significantly stronger functional connectivity (p < 0.05, FDR-corrected) with either the FFA seed or the PPA seed were selected as target ROIs for all subsequent analyses. This procedure identified 56 target ROIs in total across both hemispheres (detailed in Table 1), categorized based on their predominant functional link to either the face or scene processing networks.

### Connectome-based predictive modelling

To assess whether individual differences in resting-state functional connectivity could predict behavioral performance, we employed CPM analysis. CPM is a data-driven, cross-validated framework (Shen et al., 2017) that moves beyond simple correlation by building and rigorously testing models that predict individual behavioral scores from brain connectivity patterns. This approach provides a robust assessment of brain-behavior relationships while mitigating the risk of overfitting common in non-cross-validated analyses (Finn et al., 2015; Rosenberg & Finn, 2022; Shen et al., 2017).

We utilized the independent cohort of 371 participants described previously. Prior to modeling, specific exclusion criteria were applied. Eighteen subjects were excluded due to excessive head motion (mean framewise displacement > 0.15 mm). Subjects missing behavioral data for a specific task were further excluded from the corresponding model, resulting in final sample sizes of 350 subjects for the face-matching task model and 352 subjects for the 0-back and 2-back scene task models. Additionally, we confirmed that residual head motion was not significantly correlated with the behavioral scores within each final sample (Spearman’s rank correlation: Face-matching task: r = 0.06, p = 0.26; 0-back scene task: r = 0.03, p = 0.51; 2-back scene task: r = 0.02, p = 0.77), ensuring that prediction performance was not driven by motion artifacts.

Separate CPM analyses were performed using network-specific connectivity matrices as input. We defined two distinct networks based on the target ROI selection in Table 1. The first, a face processing network, comprised 37 regions total. This included the bilateral FFA seed regions plus the 35 ROI instances across both hemispheres identified in Table 1 as predominantly connected to the FFA seed. The second, a scene processing network, comprised 23 regions total. This included the bilateral PPA seed regions plus the 21 ROI instances across both hemispheres identified in Table 1 as predominantly connected to the PPA seed. The input connectivity matrices for each subject were thus either 37x37 (face-network) or 23x23 (scene-network), containing the fMRI partial correlations between the respective pairs of regions within that network. We used CPM to predict face-matching RTs from the face processing network connectivity and to predict 0-back and 2-back scene RTs from the scene processing network connectivity. To test the specificity of these network-behavior relationships, we also performed control analyses, predicting face-matching RTs from the scene network and predicting both scene task RTs from the face network.

For each distinct CPM analysis, a leave-one-out (LOO) cross-validation procedure was implemented across the respective final sample size. This involved iteratively training the model on N-1 subjects and testing its predictive performance on the single held-out subject, ensuring the independence between training and testing data at each fold. Within each training fold, several steps were performed following Shen et al. (2017). First, for feature selection, the strength of each edge (fMRI partial correlation value) within the relevant network matrix was correlated with the corresponding behavioral measure across the N-1 training subjects using Spearman’s rank correlation. Edges exhibiting a significant negative correlation with the behavioral score (RTs in this case; p < 0.05) were selected as predictive features. We focused specifically on negatively correlated edges as preliminary analyses indicated that positively correlated edges did not yield significant predictive models. Next, for feature summarization, a single summary score was calculated for each subject in the training set by summing the fMRI partial correlation values of all edges selected in the previous step. Finally, for model building, a simple linear regression model (y = mx + b) was fitted to the training data, predicting the behavioral scores (y) from the calculated summary scores (x). The model built in each training fold (i.e., the fitted slope ’m’ and intercept ’b’) was then applied to the left-out test subject. First, the summary score for the test subject was calculated using the same feature mask (set of predictive edges) defined from the training fold. Then, this summary score was input into the linear model derived from the training fold to generate the predicted behavioral score for that test subject. After completing all LOO folds for a given analysis, overall model performance was evaluated by calculating the Spearman’s rank correlation (r) between the predicted scores and the true observed scores across all subjects.

### Statistical analysis

All statistical analyses were performed using the Statistics and Machine Learning Toolbox in MATLAB R2023a. For the fMRI and MEG connectivity analyses comparing connectivity strengths, we used non-parametric Wilcoxon signed-rank tests. The significance threshold (alpha level) was set at p < 0.05.

Corrections for multiple comparisons across parcels were applied using the false discovery rate (FDR) procedure (Benjamini & Hochberg, 1995), with a corrected significance threshold of q < 0.05. For statistically significant target regions, z-scores were calculated from the adjusted p-values to quantify differences in connectivity strength. For the CPM analysis, the statistical significance of the prediction accuracy (correlation between predicted and observed scores) for each model was assessed non-parametrically using permutation testing (1000 iterations). In each permutation, the behavioral scores were randomly shuffled across subjects, the entire LOO CPM procedure was repeated using the shuffled scores, and the resulting correlation between predicted and shuffled scores was stored. This process built an empirical null distribution of correlation coefficients expected by chance. The final p-value for the actual model performance was calculated as the proportion of permutations that yielded a correlation coefficient greater than or equal to the correlation obtained with the true, unshuffled behavioral scores.

## Data and code availability

The data for the current study were sourced from the 1200 Subjects Release of the Human Connectome Project (Larson-Prior et al., 2013; Van Essen et al., 2013). These data are publicly accessible via ConnectomeDB (db.humanconnectome.org; (Hodge et al., 2016)).

## Acknowledgments

This work was supported by the National Science Foundation (NSF NRT NeuralStorm) under Grant No. 2152260 (to O.S.).

## Author information

### Contributions

Study conceptualization: D.B., A.S., and O.S.; Formal analysis: O.S. and A.S.; Visualization: O.S.; Writing – original draft: O.S. and A.S.; Writing – review & editing: D.B. and O.S.; Funding acquisition: D.B. and O.S.

### Corresponding author

Correspondence to Daniel Baldauf

