## Supplementary information for "Distinct resting-state connectomes for face and scene perception predict individual task performance"

### Figures

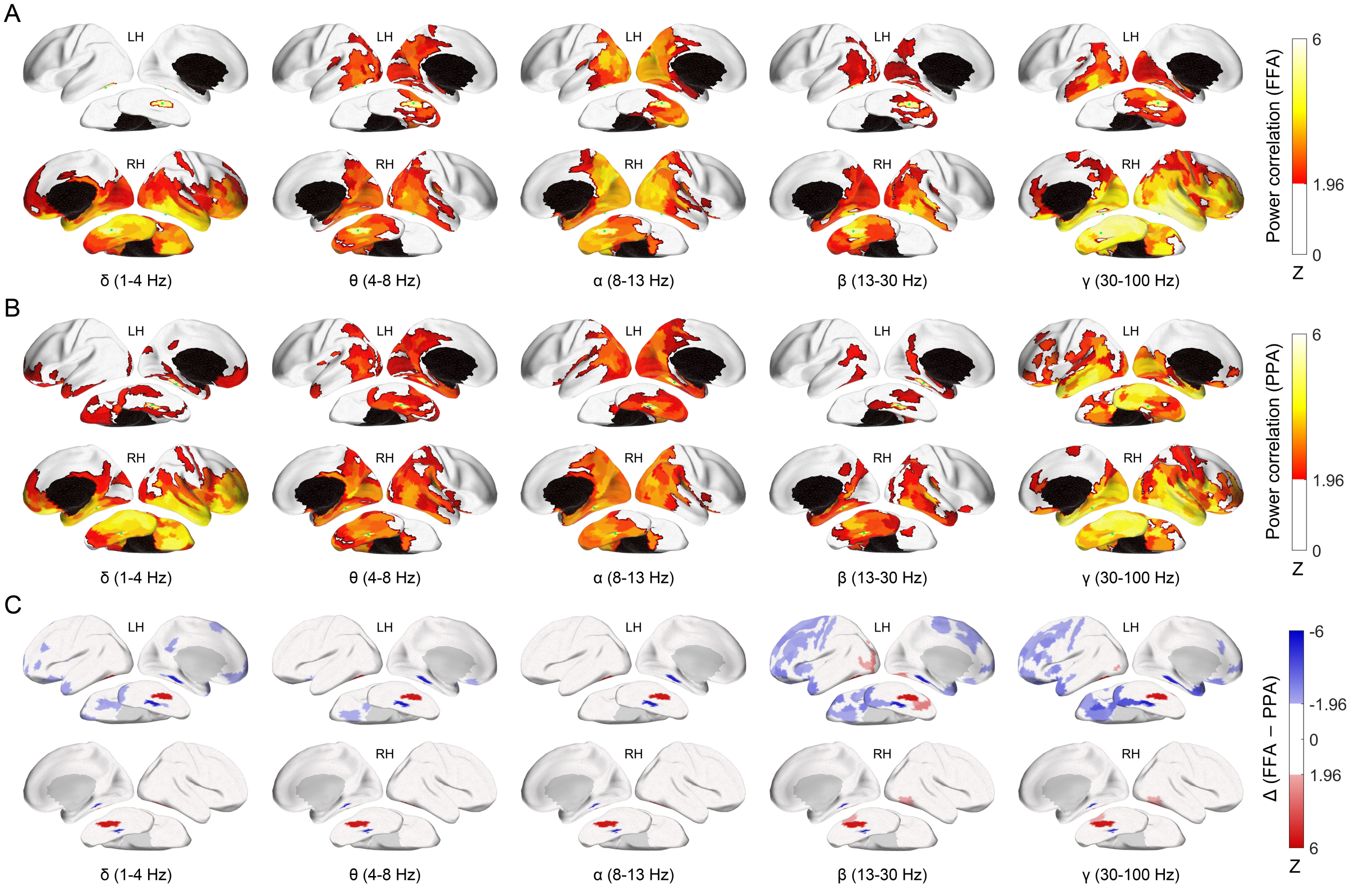


**Figure S1 | MEG power envelope correlation maps for the Fusiform Face Area (FFA) and Parahippocampal Place Area (PPA).**

(A) Whole-brain frequency-specific functional connectivity maps for the FFA based on the orthogonalized power envelope correlation method. The green dot indicates the FFA seed region. Functional connectivity is displayed for the left (LH) and right (RH) hemispheres on each ipsilateral side. (B) Same as (A) but for the PPA. (C) Direct comparison of whole-brain functional connectivity between the FFA (red) and PPA (blue), highlighting areas with preferential connectivity to each region. Statistical significance was assessed using a $p < 0.05$ threshold, FDR-corrected for 180 ROIs, and represented as z-scores.

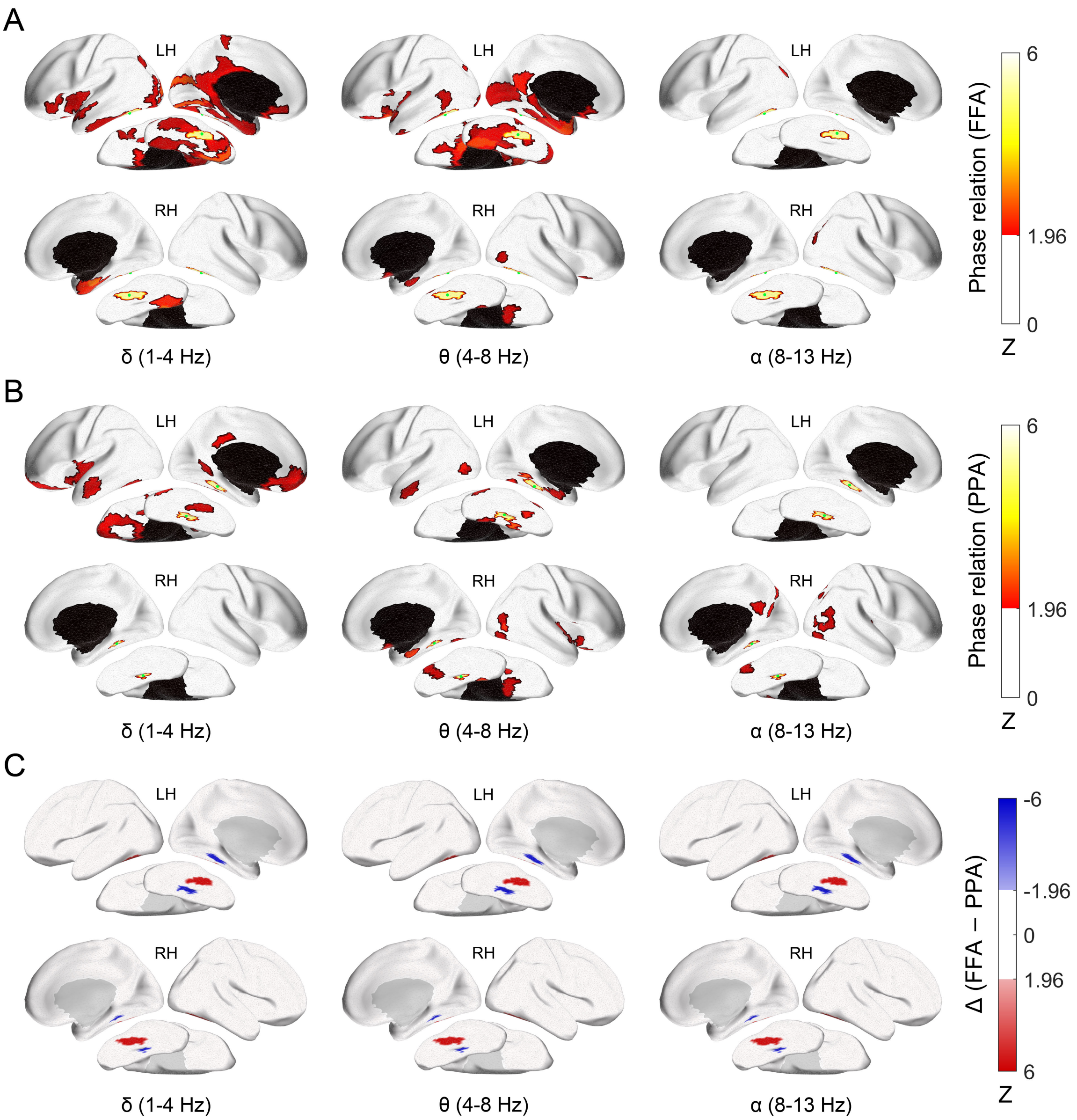


**Figure S2 | MEG phase connectivity maps for the Fusiform Face Area (FFA) and Parahippocampal Place Area (PPA).**

(A) Whole-brain frequency-specific functional connectivity maps for the FFA based on the imaginary part of the coherency method. The green dot indicates the FFA seed region. Functional connectivity is displayed for the left (LH) and right (RH) hemispheres on each ipsilateral side. (B) Same as (A) but for the PPA. There are no significant regions in beta and gamma frequency bands. (C) Direct comparison of whole-brain functional connectivity between the FFA (red) and PPA (blue), highlighting areas with preferential connectivity to each region. There were no significant areas that have predominant connectivity with either FFA or PPA across all frequency bands. Statistical significance was assessed using a $p < 0.05$ threshold, FDR-corrected for 180 ROIs, and represented as z-scores.

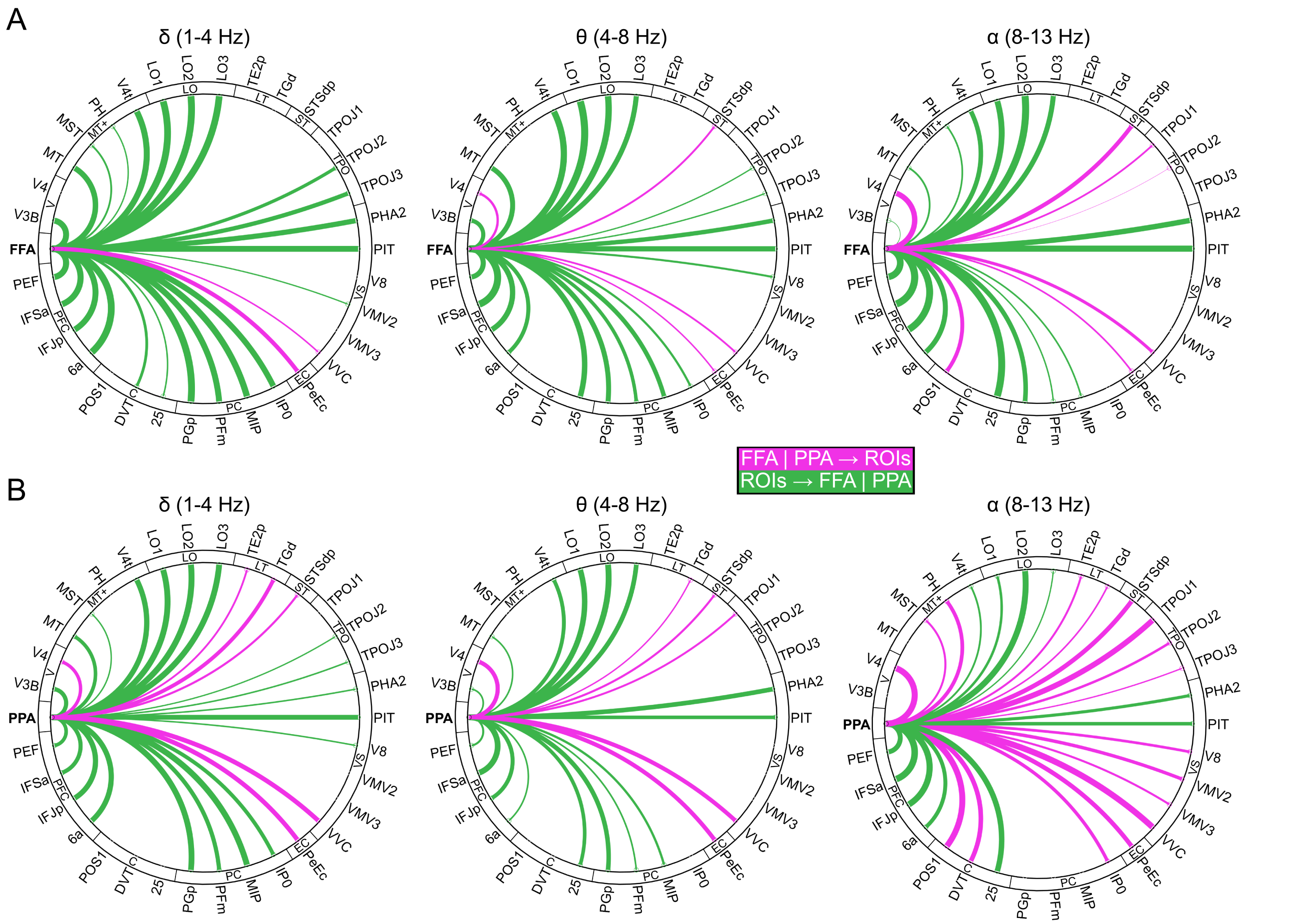


**Figure S3 | Frequency-specific directional connectivity analyses of the Fusiform Face Area (FFA) and Parahippocampal Place Area (PPA).**

(A) Circular graphs show the directional connectivity between the FFA and the brain regions listed in Table 1 across both hemispheres. Magenta lines indicate the directions of interaction from the FFA to the regions of interest (ROIs), while green lines indicate the directions of interaction from the ROIs to the FFA. (B) Same as (A) but for the PPA. Statistical significance was assessed using a $p < 0.05$ threshold, FDR-corrected for 33 ROIs, and represented as z-scores. The width of the lines reflects the strength of the connectivity. Abbreviations for the ROI groupings shown in the figure include: V = Visual areas, MT+ = Middle temporal complex, LO = Lateral occipital areas, LT = Lateral temporal cortex, ST = Superior temporal sulcus, TPO = Temporoparietal junction, VS = Ventral visual stream, EC = Entorhinal cortex, PC = Parietal cortex, C = Cingulate cortex, and PFC = Prefrontal Cortex.

### Tables

| Volume | Peak | | | Center of Mass | |
|---|---|---|---|---|---|
| (voxels) | (z-score) | XYZ (MNI) | HCP-MMP1 | XYZ (MNI) | HCP-MMP1 |
| **‘Face’ association test** | | | | | |
| 4706 | 21.0 | +40.0 -50.0 -20.0 | Right FFC | +42.3 -59.7 -13.3 | Right PH |
| 3217 | 17.4 | -40.0 -52.0 -22.0 | Left FFC | -37.8 -65.5 -13.7 | Left FFC |
| **‘Place’ association test** | | | | | |
| 481 | 11.0 | -28.0 -46.0 -10.0 | Left PHA3 | -29.7 -45.6 -10.3 | Left PHA3 |
| 361 | 8.0 | +28.0 -44.0 -14.0 | Right VVC | +29.7 -43.0 -11.7 | Right PHA3 |

**Table S1 | Principal clusters from ‘Face’ and ‘Place’ Neurosynth association test maps.**

This table summarizes the two largest activation clusters derived from Neurosynth term-based meta-analytic association maps for the terms ‘Face’ and ‘Place’ (input maps were thresholded at a False Discovery Rate of $q < 0.01$). For each reported cluster, the table lists its volume (in voxels), the peak z-score, the MNI coordinates (X,Y,Z) of both the peak activation and the center of mass, and their respective anatomical localizations within the Human Connectome Project Multi-modal Parcellation (HCP-MMP1) atlas. MNI coordinates are reported in LPI order.

| | | Left Hemisphere | | Right Hemisphere | |
|---|---|---|---|---|---|
| **Parcellation** | **Brain Region** | **P-value** | **Z-score** | **P-value** | **Z-score** |
| A4 | Auditory Cortex | <0.01 | 2.37 | – | – |
| FST | Fundus of Superior Temporal (MT+ Complex) | – | – | <0.05 | 2.05 |
| LIPd | Lateral Intraparietal Area | <0.05 | 2.07 | – | – |
| LO1 | Lateral Occipital Area | <0.01 | 2.84 | <0.001 | 3.79 |
| LO2 | Lateral Occipital Area | <0.001 | 5.12 | <0.001 | 4.82 |
| LO3 | Lateral Occipital Area | <0.001 | 4.39 | <0.001 | 4.68 |
| MST | Medial Superior Temporal (MT+ Complex) | <0.001 | 4.03 | <0.001 | 3.80 |
| MT | Middle Temporal (MT+ Complex) | <0.001 | 5.70 | <0.001 | 4.12 |
| PEF | Premotor Eye Field | – | – | <0.01 | 2.35 |
| PH | Area PH (MT+ Complex) | <0.001 | 5.90 | <0.001 | 5.91 |
| PIT | Posterior Inferior Temporal Complex | <0.001 | 5.90 | <0.001 | 5.91 |
| PeEc | Perientorhinal and Ectorhinal Complex | <0.01 | 2.78 | <0.01 | 2.73 |
| ProS | ProStriate Area | – | – | <0.01 | 2.35 |
| STSva | Superior Temporal Sulcus | <0.01 | 2.42 | – | – |
| TE1m | Middle Temporal Cortex | – | – | <0.01 | 2.43 |
| TE2p | Inferior Temporal (IT) Cortex | <0.001 | 5.90 | <0.001 | 5.91 |
| TF | Inferior Temporal (IT) Cortex | <0.001 | 5.59 | <0.001 | 5.79 |
| TGd | Temporal Polar Cortex | <0.001 | 3.26 | – | – |
| TPOJ1 | Temporo-Parieto-Occipital Junction | <0.001 | 3.54 | <0.001 | 5.02 |
| TPOJ2 | Temporo-Parieto-Occipital Junction | <0.001 | 4.96 | <0.001 | 5.10 |
| TPOJ3 | Temporo-Parieto-Occipital Junction | <0.001 | 4.85 | <0.001 | 5.07 |
| V4 | Visual Area V4 | <0.001 | 3.54 | <0.01 | 2.44 |
| V4t | Area V4t (MT+ Complex) | <0.001 | 5.55 | <0.001 | 5.79 |
| V8 | Visual Area V8 (Part of Ventral Stream) | <0.001 | 5.90 | <0.001 | 5.91 |
| VMV2 | Ventral Medial Visual Area | – | – | <0.05 | 2.08 |
| VVC | Ventral Visual Complex | <0.001 | 5.90 | <0.001 | 5.91 |

**Table S2 | Brain regions exhibiting significant functional connectivity with the Fusiform Face Area (FFA).**

The table lists areas from the Human Connectome Project multi-modal parcellation (HCP-MMP1) atlas that have significant functional coupling with the FFA ('Parcellation' column; Figure 1B). It also provides an approximate functional label for each region based on Glasser et al. (2013) ('Brain Region' column). Columns display the statistical significance (P-value) and strength (Z-score) of the functional connectivity, presented separately for the left (LH) and right (RH) hemispheres. Dashes (–) indicate that significant connectivity was not detected for that region in the specified hemisphere.

| | | Left Hemisphere | | Right Hemisphere | |
|---|---|---|---|---|---|
| **Parcellation** | **Brain Region** | **P-value** | **Z-score** | **P-value** | **Z-score** |
| 6a | Superior Premotor Cortex | – | – | <0.05 | 2.18 |
| DVT | Posterior Cingulate Cortex | <0.001 | 4.93 | <0.001 | 4.95 |
| H | Part of Hippocampus | <0.05 | 1.74 | – | – |
| IP0 | Inferior Parietal Cortex | <0.001 | 4.27 | <0.001 | 3.63 |
| PGp | Inferior Parietal Cortex | <0.001 | 5.76 | <0.001 | 5.64 |
| PH | Area PH (MT+ Complex) | <0.001 | 5.45 | – | – |
| PHA2 | Parahippocampal Area | <0.001 | 5.83 | <0.001 | 5.87 |
| POS1 | Posterior Cingulate Cortex | <0.01 | 2.63 | <0.001 | 4.46 |
| PeEc | Perientorhinal and Ectorhinal Complex | <0.001 | 5.80 | <0.001 | 3.15 |
| TE2p | Lateral Temporal Cortex | <0.05 | 2.05 | – | – |
| TF | Lateral Temporal Cortex | <0.001 | 5.24 | <0.001 | 5.87 |
| VMV2 | Ventral Medial Visual Area | <0.001 | 5.83 | <0.001 | 5.87 |
| VMV3 | Ventral Medial Visual Area | <0.001 | 4.58 | <0.001 | 4.33 |
| VVC | Ventral Visual Complex | <0.001 | 5.83 | <0.001 | 5.87 |

**Table S3 | Brain regions exhibiting significant functional connectivity with the Parahippocampal Place Area (PPA).**

The table lists areas from the Human Connectome Project multi-modal parcellation (HCP-MMP1) atlas that have significant functional coupling with the PPA ('Parcellation' column; Figure 1C). It also provides an approximate functional label for each region based on Glasser et al. (2013) ('Brain Region' column). Columns display the statistical significance (P-value) and strength (Z-score) of the functional connectivity, presented separately for the left (LH) and right (RH) hemispheres. Dashes (–) indicate that significant connectivity was not detected for that region in the specified hemisphere.
